# Regulated RIAM–Talin Engagement Controls Adhesion Stability and Mechanical Output

**DOI:** 10.64898/2026.01.15.699718

**Authors:** Nikhil Mittal, Ho-Sup Lee, Mark H. Ginsberg, Sangyoon J. Han

**Affiliations:** Department of Biomedical Engineering, Michigan Technological University, Houghton, MI, USA; Health Research Institute, Michigan Technological University, Houghton, MI, USA; University of California San Diego School of Medicine, La Jolla, CA, USA; Department of Mechanical Engineering and Engineering Mechanics, Michigan Technological University, Houghton, MI, USA

**Keywords:** Focal adhesion, RIAM, talin-1, Rap1, phosphoinositides, traction force

## Abstract

RIAM (Rap1-GTP–interacting adaptor molecule) links Rap1 to talin-1 to promote integrin activation and is normally enriched at nascent adhesions. How the duration of RIAM–talin engagement influences adhesion dynamics and force transmission remains unclear. Here, we used a RIAM chimera in which the native talin-binding site was replaced with the talin-binding motif of Kank2 to enforce sustained talin association and redistribute RIAM modules to mature adhesions. This talin-tethered RIAM chimera enhanced integrin activation, accelerated both adhesion assembly and disassembly, reduced adhesion lifetime, increased the fraction of nascent adhesions nucleated by RIAM, and elevated cellular traction forces. Notably, while integrin activation by the chimera was largely Rap1-independent, Rap1 binding remained necessary for optimal adhesion turnover kinetics and full traction force generation. These findings demonstrate that prolonged talin engagement of RIAM is sufficient to reprogram adhesion dynamics and reveal a separable role for Rap1 in coordinating force transmission downstream of integrin activation. Together, our results highlight the importance of regulating the residence and engagement mode of RIAM at talin for controlling adhesion plasticity and mechanotransduction.

## Introduction

Cell adhesion to the extracellular matrix (ECM) is a crucial factor for various cellular behaviors, influencing processes such as angiogenesis, inflammation, tumor progression, and vascular development (1–3). These functions are largely mediated by integrins, transmembrane receptors that connect extracellular ligands to the actin cytoskeleton and transmit mechanical and chemical signals bidirectionally (4). Through this linkage, integrins support bidirectional signaling that enables cells to sense and respond to mechanical cues in their environment. A central regulator of integrin activation is talin-1, a large cytoskeletal adaptor that binds β-integrin cytoplasmic tails and connects them to filamentous actin, thereby stabilizing integrins in a high-affinity ligand-binding conformation (5–7).

One major pathway for talin recruitment is mediated by RIAM (RAP1-GTP-interacting adaptor molecule), which is a member of the MRL (Mig10–RIAM–Lamellipodin) adapter protein family (8). RIAM contains multiple functional domains including an N-terminal region harboring two talin-binding sites (TBS1 and TBS2), a Ras association (RA) domain, a pleckstrin homology (PH) domain, and proline-rich motifs (9). Among these, TBS1 is the primary determinant of talin engagement and integrin activation, whereas TBS2 exhibits weaker talin affinity and limited activating capacity (9–11). In cells, RIAM is enriched at nascent adhesions (NAs) near the leading edge, where integrins are initially activated (12) but is comparatively sparse in mature focal adhesions (FAs), which are dominated by force-bearing components such as vinculin (13). Recent structural and functional studies have suggested potential intramolecular interactions between RIAM domains (14), but the regulatory impact of these interactions on talin-1 binding, integrin activation, and downstream mechanical signaling has not been clearly elucidated. Moreover, while RIAM is known to support integrin activation, its direct role in adhesion turnover and traction force generation remains unclear—particularly whether disrupting RIAM’s autoinhibition could modulate its function beyond adhesion recruitment.

Recent structural studies have revealed that RIAM can adopt an autoinhibited conformation in which its talin-binding activity is masked by intramolecular interactions involving the PH domain and upstream inhibitory segments (15). In this state, RIAM is prevented from prematurely engaging talin in the cytosol. Binding of Rap1-GTP to the RA domain, together with membrane recruitment through phosphoinositides and phosphorylation-dependent regulation, can relieve this autoinhibition and promote RIAM-dependent integrin activation at the plasma membrane (16). While this regulatory mechanism is well established for integrin activation, far less is known about how the *mode and duration* of RIAM–talin engagement influence downstream adhesion dynamics, turnover, and force transmission once adhesions experience mechanical load.

Importantly, adhesion maturation is not solely determined by integrin activation, but also by the coordinated assembly and disassembly of adhesion complexes under force. Nascent adhesions can either disassemble rapidly or mature into force-transmitting focal adhesions, a process that depends on talin engagement, actomyosin contractility, and vinculin recruitment (10,13). Given RIAM’s transient enrichment at early adhesions, a key unanswered question is whether limiting RIAM–talin engagement is necessary for proper adhesion maturation, or whether prolonged RIAM association with talin can actively reprogram adhesion mechanics and turnover.

To address this question, we employed a synthetic perturbation strategy that enforces sustained talin engagement by RIAM. Specifically, we engineered a RIAM chimera in which the native talin-binding site (TBS1) was replaced with the high-affinity talin-binding motif of Kank2, a focal adhesion–associated protein that binds the talin R7 domain without undergoing intramolecular autoinhibition. This design does not aim to recapitulate native RIAM activation, but rather to prolong RIAM’s association with talin and redistribute its functional modules within adhesion complexes. Using this talin-tethered RIAM chimera, we examined how sustained RIAM–talin engagement influences integrin activation, adhesion assembly and disassembly kinetics, adhesion lifetime, and traction force generation. Our results show that enforced talin engagement of RIAM enhances integrin activation, accelerates both adhesion assembly and disassembly, increases the fraction of nascent adhesions nucleated by RIAM, and elevates traction forces in a partially Rap1-dependent manner. These findings reveal that the temporal regulation of RIAM–talin interaction is a critical determinant of adhesion plasticity and force transmission, and establish prolonged RIAM engagement as a potent modulator of mechanosensitive adhesion dynamics.

## Results

### A talin-tethered RIAM chimera bypasses PH-mediated sequestration and activates integrins independently of Rap1

RIAM plays a critical role in integrin activation and adhesion assembly (17), but its activity is tightly regulated through autoinhibition involving its NH₂-terminal talin-binding site (TBS1) and the adjacent PH domain (15, 18). Structural and biochemical studies have shown that these interactions limit talin engagement unless RIAM is recruited to the plasma membrane by Rap1-GTP and phosphoinositide (15). While this regulatory mechanism is well established, the consequences of enforcing sustained talin engagement by RIAM in cells remain unclear.

To experimentally enforce talin engagement while avoiding RIAM’s native intramolecular sequestration, we engineered a chimeric RIAM construct in which the native TBS1 (residues 1–30) was replaced by the talin-binding motif of Kank2 (residues 31–65). The Kank2 talin-binding sequence binds the folded talin R7 domain with high affinity and does not require unfolding of the talin rod (19), making it an effective tool to tether RIAM to talin independently of RIAM’s endogenous regulatory interfaces.

We first tested whether the Kank2 talin-binding motif can substitute for RIAM TBS1 in binding talin. Co-immunoprecipitation assays in CHO cells revealed that both GFP-tagged RIAM(1–30) and the GFP-tagged Kank2(31–65) motif efficiently associated with full-length FLAG-talin-1 (Fig. 1A), confirming that the heterologous Kank2 sequence retains robust talin-binding capacity. We next examined whether this talin-binding motif is subject to sequestration by the RIAM PH domain. As expected, GFP-RIAM(1–30) strongly associated with the FLAG-tagged RIAM PH domain, consistent with previously described intramolecular interactions. In contrast, GFP-Kank2(31–65) failed to bind the RIAM PH domain (Fig. 1B), indicating that the Kank2 talin-binding motif lacks the interface required for PH-mediated sequestration. These results establish that the Kank2 talin-binding motif can engage talin while remaining uncoupled from RIAM’s PH-dependent intramolecular interactions.

**Figure 1.**
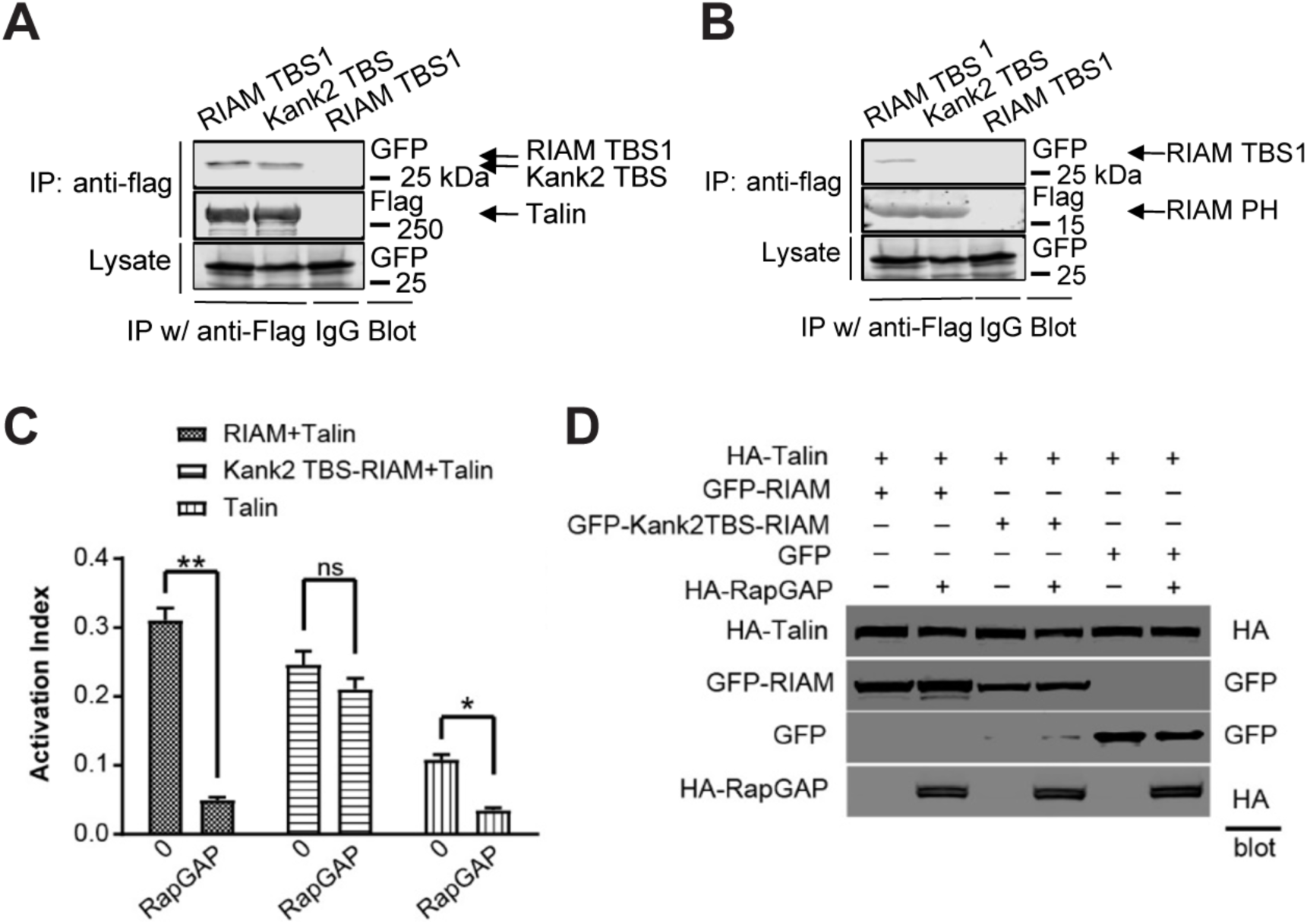
Talin binding and Rap1-independent integrin activation by Kank2TBS-RIAM. (A and B) Kank2TBS (31-65) binds talin-1, but not RIAM PH domain. (A) Co-immunoprecipitation of FLAG-talin-1 with GFP-tagged RIAM TBS1 (1–30) or GFP-Kank2TBS (31–65); detection by anti-FLAG and anti-GFP immunoblotting. (B) Co-immunoprecipitation of FLAG-RIAM PH with GFP-tagged RIAM TBS1 (1–30) or GFP-Kank2TBS (31–65); detection by anti-FLAG and anti-GFP immunoblotting. (C–D) Kank2TBS-RIAM activates integrins independently of Rap1-mediated regulation. (C) Flow-cytometric analysis of αIIbβ3 integrin activation in CHO cells co-transfected with HA-talin and either GFP-RIAM, GFP-Kank2TBS-RIAM, or GFP control, in the presence or absence of HA-RapGAP. Activation index derived from PAC-1 antibody binding. (D) Immunoblot validation of protein expression in samples from (C) using anti-HA and anti-GFP antibodies.

We next asked how enforced talin engagement affects integrin activation and whether this process remains dependent on Rap1. CHO cells expressing αIIbβ3 integrins were co-transfected with talin and either GFP-RIAM or GFP-Kank2TBS-RIAM, in the presence or absence of RapGAP to suppress Rap1 activity. Integrin activation was quantified by flow cytometry using binding of the activation-specific antibody PAC-1 (20). Consistent with previous studies, RIAM-mediated integrin activation was strongly Rap1-dependent, as RapGAP expression markedly reduced PAC-1 binding (Fig. 1C, left). In contrast, the Kank2TBS-RIAM chimera induced robust integrin activation even in the presence of RapGAP (Fig. 1C, middle), indicating that enforced talin engagement bypasses the requirement for Rap1-mediated RIAM activation. Talin expressed alone produced moderate integrin activation that was partially sensitive to RapGAP (Fig. 1C, right), likely reflecting residual endogenous RIAM activity. Immunoblotting confirmed comparable expression of HA-talin, GFP-RIAM constructs, and HA-RapGAP across all conditions (Fig. 1D). Together, these data demonstrate that tethering RIAM to talin via the Kank2 talin-binding motif is sufficient to activate integrins independently of Rap1. This chimera therefore provides a tool to examine how prolonged talin engagement by RIAM—decoupled from its native regulatory constraints—affects adhesion dynamics and force transmission downstream of integrin activation.

### Enforced talin engagement enhances RIAM recruitment but does not bypass Rap1-dependent adhesion assembly

To examine how enforced talin engagement affects RIAM localization and adhesion dynamics independently of Rap1 binding, we generated a Rap1-binding–deficient variant of the chimera, Kank2TBS-RIAM(K210E), in which a point mutation in the RA domain disrupts Rap1 interaction. Inducible GFP-tagged RIAM, Kank2TBS-RIAM, and Kank2TBS-RIAM(K210E) were expressed in 3T3 fibroblasts stably expressing paxillin-mRuby, followed by imaging using total internal reflection fluorescence (TIRF) microscopy.

Compared with GFP-RIAM, which exhibited predominantly diffuse cytoplasmic localization with minimal enrichment at paxillin-positive adhesions (Fig. S1A–D), GFP-Kank2TBS-RIAM showed robust accumulation at adhesions throughout the ventral surface (Fig. S1E–H). The Rap1-binding–deficient K210E mutant also localized to adhesions, although with reduced enrichment relative to the active chimera (Fig. S1I–L). Quantitative analysis across nascent, focal, and fibrillar adhesion populations confirmed that Kank2TBS-RIAM significantly increased RIAM enrichment at adhesions, whereas the K210E mutation partially attenuated this effect (Fig. S1M–O). In addition, expression of Kank2TBS-RIAM was associated with the formation of elongated, fibrillar adhesion structures (Fig. S1P–R), indicating that sustained RIAM–talin engagement alters adhesion architecture.

### Kank2TBS-RIAM Enhances Early Adhesion Recruitment but Requires Rap1 for Efficient Assembly

We next asked how these changes in RIAM localization influence adhesion assembly dynamics. Live-cell TIRF imaging was used to track individual nascent adhesions over time using the paxillin channel, while simultaneously monitoring the recruitment kinetics of the GFP-tagged RIAM constructs. In cells expressing GFP-RIAM, GFP fluorescence remained diffuse and showed weak temporal coupling to paxillin accumulation during adhesion assembly (Fig. 2A–D). In contrast, GFP-Kank2TBS-RIAM was recruited earlier and more strongly to assembling adhesions, closely tracking paxillin intensity (Fig. 2E–H). The K210E mutant exhibited intermediate recruitment behavior, with enhanced adhesion association compared to GFP-RIAM but reduced enrichment relative to the active chimera (Fig. 2I–L).

**Figure 2.**
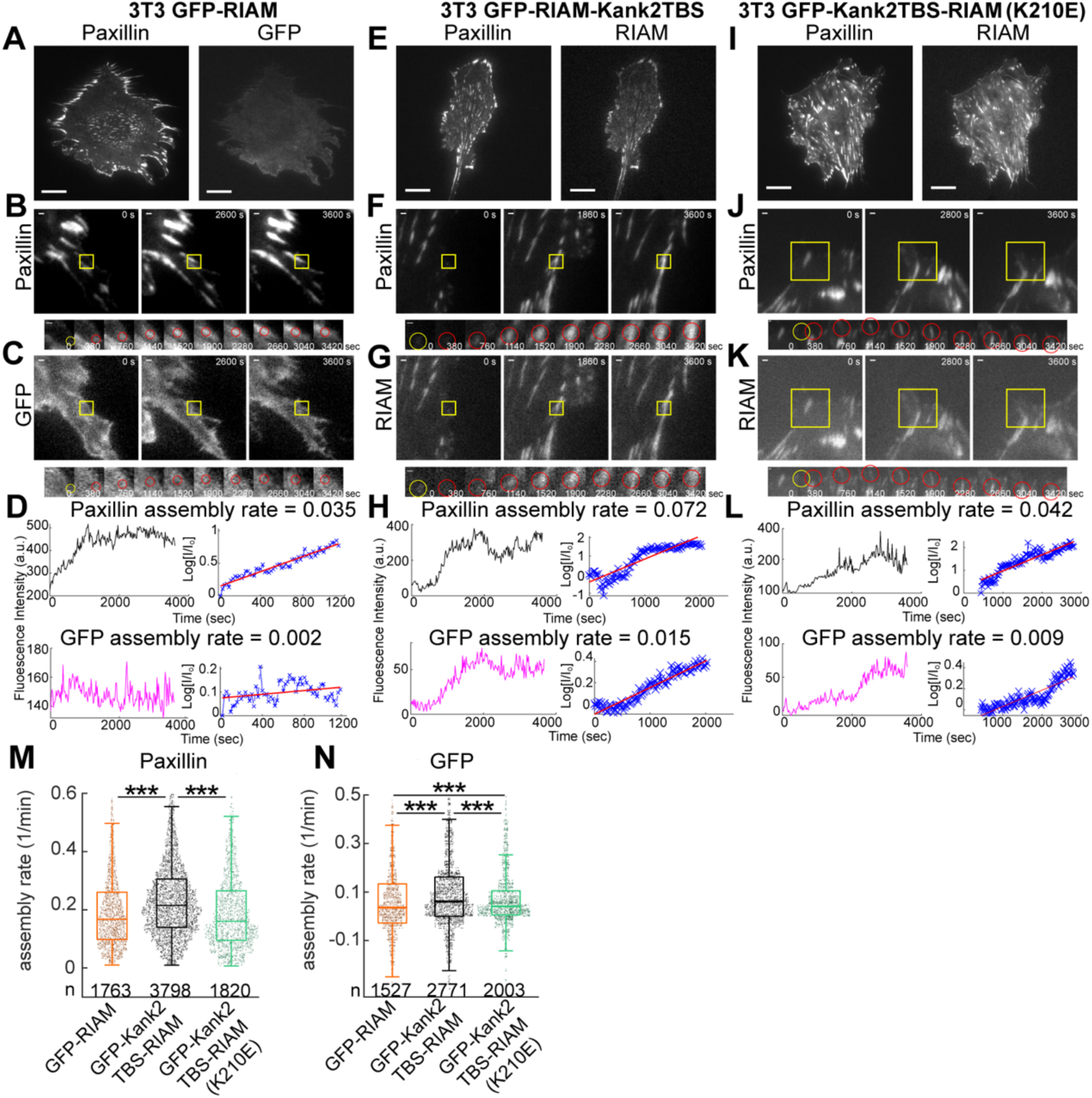
RIAM-Kank2TBS promotes faster assembly of NAs. (A) Representative images of paxillin-mRuby (left) and GFP-RIAM (right) in 3T3 cells co-expressing paxillin-mRuby and GFP-RIAM. (B, C) Representative intensity traces of paxillin (B) and GFP (C) during adhesion assembly. (D) Fluorescence intensity and assembly kinetics of paxillin (top) and GFP (bottom). (E) Representative images of paxillin-mRuby (left) and GFP (right) in cells expressing GFP-Kank2TBS-RIAM. (F, G) Representative traces of paxillin (F) and GFP-RIAM (G) during adhesion assembly. (H) Fluorescence intensity and assembly kinetics of paxillin (top) and GFP (bottom). (I) Representative images of paxillin-mRuby (left) and GFP (right) in cells expressing GFP-Kank2TBS-RIAM(K210E). Scale bar: 20 μm (A, E, I). (J, K) Representative traces of paxillin (J) and GFP (K) during adhesion assembly. (L) Fluorescence intensity and assembly kinetics of paxillin (top) and GFP (bottom). Yellow boxes indicate regions of tracked adhesions. Scale bar: 1 μm. I₀ = initial fluorescence intensity, I = intensity at the indicated time points. Rate constants for assembly determined from slopes of log(I/I₀) plots. (M) Boxplot of paxillin assembly rates for 3T3 cells expressing GFP-RIAM, GFP-Kank2TBS-RIAM, and GFP-Kank2TBS-RIAM(K210E). (N) Boxplot of GFP assembly rates for the same conditions: GFP-RIAM, GFP-Kank2TBS-RIAM, GFP-Kank2TBS-RIAM(K210E). Data collected from five cells per condition. **p < 1 × 10⁻²; ***p < 1 × 10⁻⁵ (Mann–Whitney U test).

Despite these differences in RIAM recruitment, quantitative analysis of paxillin assembly kinetics revealed no acceleration of adhesion assembly beyond the level observed in GFP-RIAM–expressing cells. While Kank2TBS-RIAM significantly increased the assembly rate of the GFP signal, paxillin assembly rates were unchanged relative to the RIAM control (Fig. 2M,N). Notably, disruption of Rap1 binding in the K210E mutant fully restored paxillin assembly rates to control levels while only partially restoring RIAM recruitment kinetics. These results indicate that enhanced RIAM localization driven by enforced talin engagement is not sufficient to accelerate adhesion assembly and that Rap1 interaction remains necessary for efficient coordination of adhesion maturation.

Together, these findings demonstrate that prolonged talin engagement uncouples RIAM localization from productive adhesion assembly. While tethering RIAM to talin promotes its recruitment to adhesions, Rap1-dependent regulation is still required to translate RIAM enrichment into efficient adhesion growth, revealing a separable role for Rap1 in organizing adhesion assembly dynamics downstream of integrin activation.

### Prolonged RIAM–talin engagement accelerates adhesion disassembly and requires Rap1 for coordinated turnover

Previous studies have shown that RIAM contributes to focal adhesion turnover downstream of integrin activation, in part through signaling pathways involving focal adhesion kinase (FAK), and that depletion of RIAM leads to enlarged, stabilized adhesions with impaired disassembly and reduced cell motility (21–23). These observations suggest that RIAM not only promotes adhesion formation but also participates in adhesion removal. We therefore asked whether enforced talin engagement via the Kank2TBS-RIAM chimera alters the kinetics or coordination of adhesion disassembly.

Using the same live-cell TIRF time-lapse datasets analyzed for assembly dynamics in Figure 2, we focused on the disassembly phase of individual paxillin-positive adhesions, quantified using the method established by Webb and colleagues (24). In cells expressing GFP-RIAM, disassembling adhesions showed minimal enrichment of GFP signal and weak temporal coupling between RIAM intensity and paxillin loss (Fig. 3A–D), consistent with the transient association of wild-type RIAM with early adhesion structures. In contrast, adhesions in cells expressing GFP-Kank2TBS-RIAM underwent more abrupt and synchronized disassembly, characterized by steeper paxillin intensity decay and tighter temporal coordination across individual events (Fig. 3E–H). Quantitative analysis confirmed a significant increase in paxillin disassembly rates relative to the GFP-RIAM condition (Fig. 3M), indicating accelerated adhesion turnover when RIAM is persistently engaged with talin.

**Figure 3.**
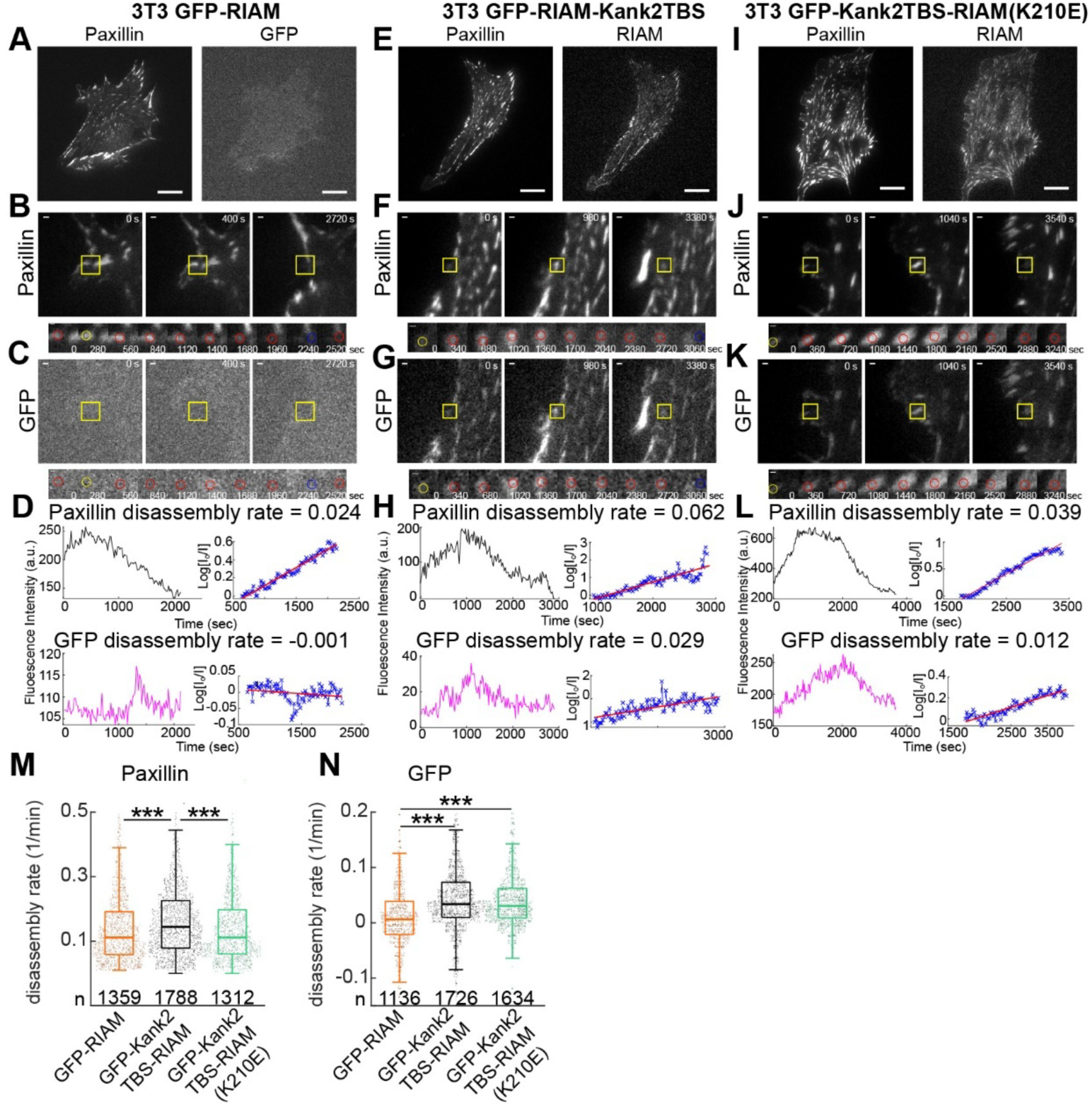
RIAM-Kank2TBS promotes faster disassembly of NAs in a Rap1-dependent manner. (A) Represent image of Paxillin-mRuby (left) and diffused GFP-RIAM (right) in 3T3 mRuby-paxillin GFP-RIAM. (B, C) Representative traces of paxillin (B) and GFP (C) of disassembling adhesions in 3T3 mRuby-paxillin GFP-RIAM. (D) Traces of fluorescence intensity (left) and the kinetics of disassembly (right) for paxillin (top) and GFP (bottom). (E) Represent image of Paxillin-mRuby (left) and GFP (right) in mRuby-paxillin GFP-RIAM Kank2TBS. (F, G) Representative traces of paxillin (F) and GFP-RIAM (G) of disassembling adhesions in 3T3 mRuby-paxillin GFP-RIAM-Kank2TBS. (H) Traces of fluorescence intensity (left) and the kinetics of disassembly (right) for paxillin (top) and GFP (bottom). (I) Represent image of Paxillin-mRuby (left) and GFP (right) in mRuby-paxillin GFP-RIAM Kank2TBS (K210E). (J, K) Representative traces of paxillin (J) and GFP (K) of disassembly adhesions in mRuby-paxillin GFP-RIAM Kank2TBS (K210E). (L) Traces of fluorescence intensity (left) and the kinetics of disassembly (right) for paxillin (top) and GFP (bottom). Scale bar: 20μm (A, E, I). Yellow boxes show positions of the example adhesions in time-lapse image sequences of mRuby-paxillin and GFP in 3T3 mRuby-paxillin and GFP-RIAM (paxillin: B, top; GFP: C, bottom), mRuby-paxillin and GFP-RIAM Kank2TBS (paxillin: F, top; GFP: G, bottom), and mRuby-paxillin and GFP-RIAM Kank2TBS (K210E) (paxillin: J, top; GFP: K, bottom). Scale bar: 1 μm. I0 is the initial fluorescent intensity, and I is the fluorescent intensity at the indicated times. Rate constants for disassembly are determined from the slopes of these graphs of Log(I0/I). (M) A boxplot of the disassembly rates of mRuby-paxillin for 3T3s expressing GFP-RIAM (n=1763), GFP-RIAM-Kank2TBS (n=3798), and mRuby-paxillin GFP-RIAM Kank2TBS (K210E) (n=1312). (N) A box plot of disassembly rates of GFP for 3T3s expressing GFP-RIAM (n=1527), GFP-RIAM-Kank2TBS (n=2771), and mRuby-paxillin GFP-RIAM Kank2TBS (K210E) (n=1634). Sample numbers extracted from 5 cells for both paxillin and GFP per condition. **p<1×10^-2^ ***, p<1×10^-5^ by Mann-Whitney U test.

The Rap1-binding–deficient Kank2TBS-RIAM(K210E) mutant localized efficiently to adhesions (Fig. 3I–K) but did not recapitulate the accelerated disassembly phenotype. Paxillin disassembly kinetics in this condition were comparable to those observed in GFP-RIAM–expressing cells (Fig. 3L, M). Notably, while paxillin loss proceeded at near-control rates, the GFP signal from the K210E mutant exhibited delayed and heterogeneous dissociation from adhesions (Fig. 3N), suggesting a decoupling between RIAM removal and adhesion disassembly.

Together, these results indicate that prolonged RIAM–talin engagement promotes rapid adhesion disassembly, but that Rap1 binding is required for efficient temporal coordination between RIAM dissociation and paxillin turnover. Thus, while Rap1 interaction is not strictly required to initiate adhesion disassembly, it appears to play a critical role in synchronizing molecular disengagement during adhesion turnover. These findings further support the idea that RIAM contributes to both assembly and disassembly phases of adhesion dynamics, with Rap1 serving as a key regulator of the timing and coordination of these processes.

### Prolonged RIAM–talin engagement shortens adhesion lifetime while increasing nascent adhesion nucleation

Rapid adhesion assembly is often accompanied by rapid disassembly, leading to a net decrease in adhesion lifetime unless counterbalanced by sustained nucleation of new adhesions to maintain adhesion density and cell–matrix connectivity (24–26). Given that the Kank2TBS-RIAM chimera accelerates both adhesion assembly and disassembly (Figs. 2 and 3), we next asked how prolonged RIAM–talin engagement influences adhesion lifetime and nascent adhesion (NA) nucleation.

We first quantified adhesion lifetimes as a measure of adhesion stability. Cells expressing GFP-Kank2TBS-RIAM exhibited significantly shorter lifetimes of both mature focal adhesions (FAs; Fig. 4A) and nascent adhesions that matured into FAs (Fig. 4B) compared to GFP-RIAM–expressing controls. These reduced lifetimes are consistent with the accelerated disassembly kinetics observed for the chimera (Fig. 3). In contrast, adhesions in cells expressing the Rap1-binding–deficient mutant Kank2TBS-RIAM(K210E) displayed significantly longer lifetimes (Fig. 4A,B), indicating that Rap1 interaction contributes to efficient adhesion turnover.

**Figure 4.**
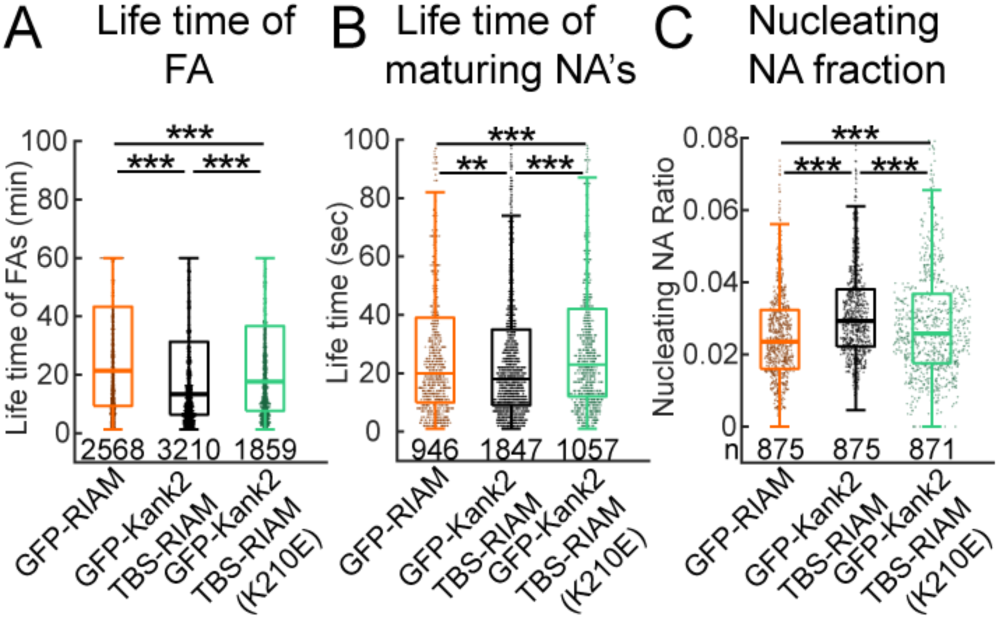
RIAM increases NA nucleation but decreases the lifetime of FA and maturing adhesions. (A) Nucleating NA for cells expressing GFP-RIAM, GFP-RIAM-Kank2TBS, and GFP-RIAM-Kank2TBS(K210E). (B) The lifetime of FAs for 3T3s expressing GFP-RIAM, GFP-RIAM-Kank2TBS, and GFP-RIAM-Kank2TBS (K210E). (C) Lifetimes of non-maturing NAs for cells expressing GFP-RIAM (n=946), GFP-RIAM-Kank2TBS, and GFP-RIAM-Kank2TBS(K210E). Sample numbers were extracted from 5 cells for both paxillin and GFP per each condition. **p<1×10^-2^ ***.: p<1×10^-5^ by Mann-Whitney U test.

We next examined whether shortened adhesion lifetimes were offset by increased adhesion initiation. Indeed, expression of GFP-Kank2TBS-RIAM significantly increased the fraction of nucleating nascent adhesions compared to GFP-RIAM (Fig. 4C), indicating enhanced initiation of productive adhesions. This effect is consistent with elevated integrin activation driven by the chimera (Fig. 1C). Importantly, the increased nucleation efficiency was attenuated in the Rap1-binding–deficient K210E mutant (Fig. 4C), suggesting that while enforced talin engagement promotes integrin activation, Rap1 signaling enhances the efficiency with which nascent adhesions are productively initiated. Together, these results indicate that prolonged RIAM–talin engagement shortens adhesion lifetime while simultaneously increasing nascent adhesion nucleation. Rap1 does not simply act as an on/off switch for RIAM activity, but instead selectively tunes the kinetics of RIAM recruitment and removal, thereby coordinating adhesion initiation and turnover. This balance enables sustained adhesion remodeling despite rapid individual adhesion turnover, providing a mechanistic link between RIAM–talin engagement, integrin activation, and dynamic adhesion plasticity.

### Prolonged RIAM–talin engagement enhances adhesion-based traction in a Rap1-dependent manner

Focal adhesion size and maturation have often been correlated with increased traction force transmission (27, 28), and rapid force growth has been linked to efficient nascent adhesion (NA) maturation and accelerated adhesion assembly (29, 30). Conversely, fast adhesion disassembly is generally thought to limit sustained force transmission. Given that the RIAM-Kank2TBS chimera accelerates both adhesion assembly and disassembly, we asked whether this altered adhesion dynamics is compatible with effective traction generation.

To address this, we performed traction force microscopy (TFM) on cells expressing GFP-RIAM, GFP-Kank2TBS-RIAM, or the Rap1-binding–deficient mutant GFP-Kank2TBS-RIAM(K210E), plated on compliant 5 kPa silicone substrates (Fig. 5A–C). Cells expressing GFP-Kank2TBS-RIAM generated robust traction forces that were broadly distributed along the cell periphery (Fig. 5B), in contrast to GFP-RIAM–expressing cells, which exhibited markedly lower traction magnitudes and more spatially restricted force patterns (Fig. 5A). In comparison, the Rap1-binding–deficient K210E mutant produced weaker and more localized traction forces (Fig. 5C).

**Figure 5.**
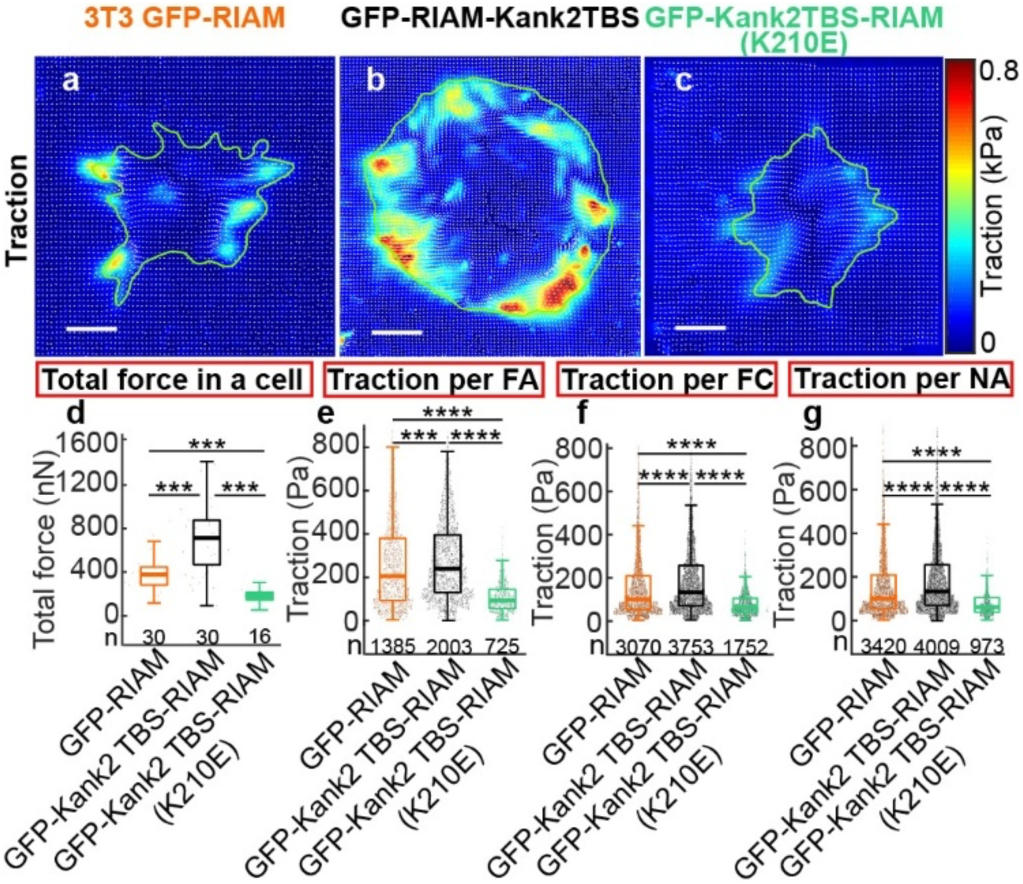
RIAM chimera promotes larger traction transmission than control in a Rap1-dependent manner. (A, B, C) RIAM chimera promotes larger traction transmission than control in a Rap1-dependent manner. High-resolution traction maps, with traction vectors (white arrows), of a representative cell expressing GFP-RIAM (A), GFP-RIAM Kank2TBS (B), and GFP-RIAM Kank2TBS(K210E) (C) on a 5kPa gel. Scale bar: 20 μm (D-G). Box plots of the total force, i.e., total traction integrated over cell area (D), average traction magnitude in FAs (E), in FCs (F), and NAs (G). The numbers indicated under each box plot represent the number of cells for (D) and the number of adhesions for (E-G). The number of adhesions was collected from multiple cells, i.e., n=30 for cells expressing GFP-RIAM, n=30 for GFP-RIAM-Kank2TBS (n=30), and n=16 for cells expressing GFP-RIAM-Kank2TBS(K210E). ***: p<1×10-5, ****: p<1×10-10 by Mann-Whitney U test.

Quantification of total cellular traction force, defined as the sum of traction magnitudes integrated over the cell area, revealed a significant increase in force generation in GFP-Kank2TBS-RIAM–expressing cells relative to GFP-RIAM controls (Fig. 5D). Importantly, this enhancement was strongly Rap1 dependent, as total traction forces in cells expressing the K210E mutant were reduced to levels below those observed in GFP-RIAM–expressing cells (Fig. 5D). Consistent trends were observed when traction forces were analyzed at the level of individual focal adhesions (FAs), focal complexes (FCs), and nascent adhesions (NAs) (Fig. 5E–G), indicating that the effect of the chimera on force transmission is not restricted to a specific adhesion class.

Together, these results demonstrate that prolonged RIAM–talin engagement can support elevated traction force generation despite rapid adhesion turnover. While enforced talin binding via the Kank2TBS promotes force transmission, Rap1 interaction remains essential for achieving maximal force output and proper spatial coordination of traction. These findings indicate that RIAM not only regulates the kinetics of adhesion assembly and disassembly but also contributes to the mechanical efficiency of force transmission, with Rap1 selectively tuning the magnitude and organization of adhesion-based forces.

## Discussion

In this study, we identify RIAM autoinhibition and its timely replacement at talin as key determinants of adhesion dynamics and mechanical output. As summarized in Fig. 6, wild-type RIAM exists in an autoinhibited conformation in which intramolecular interactions between TBS1 and the PH domain limit talin engagement. Activation by Rap1 relieves this inhibition, enabling RIAM to recruit talin to integrins and initiate nascent adhesion assembly. Under increasing mechanical load, RIAM is subsequently displaced by vinculin, which binds unfolded talin and stabilizes the adhesion, thereby supporting sustained force transmission. By contrast, replacing RIAM TBS1 with the high-affinity Kank2 talin-binding motif generates a constitutively active RIAM that bypasses autoinhibition and remains persistently associated with talin under force. This prolonged RIAM–talin interaction uncouples adhesion dynamics from the normal RIAM–vinculin exchange, resulting in enhanced adhesion assembly and turnover, shortened adhesion lifetime, and elevated traction in a Rap1-dependent manner. Together, these findings support a model in which RIAM and vinculin define sequential mechanical states of talin engagement, and in which the timely replacement of RIAM by vinculin functions as a molecular timer that limits excessive Rap1-dependent turnover while enabling stable, load-bearing adhesions.

**Figure 6.**
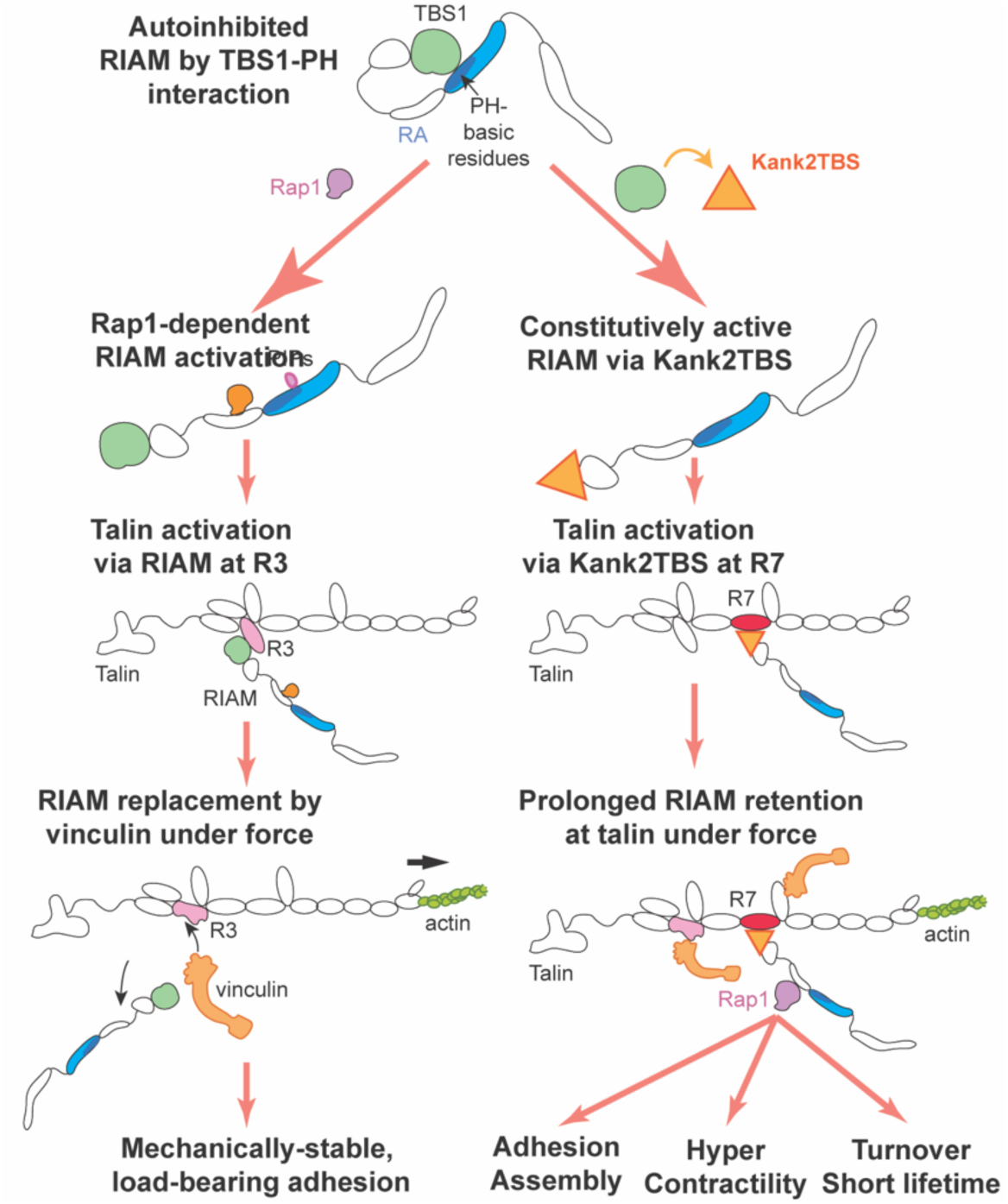
Model of RIAM autoinhibition, activation, and replacement during adhesion maturation and force transmission. Wild-type RIAM is autoinhibited through an intramolecular interaction between TBS1 and the PH domain, restricting talin binding. Rap1 activation relieves this autoinhibition, allowing RIAM to engage talin at the R2/R3 region and promote integrin activation and nascent adhesion assembly. Under mechanical load, RIAM is normally replaced by vinculin, which binds unfolded talin and stabilizes adhesions for sustained force transmission. In contrast, a RIAM–Kank2TBS chimera bypasses autoinhibition and binds talin at the R7 domain, leading to prolonged RIAM retention under force. This extended association promotes hyper-dynamic adhesions characterized by enhanced assembly, elevated traction, accelerated turnover, and shortened adhesion lifetime. The model highlights RIAM–vinculin exchange as a key regulatory step that governs the transition from dynamic adhesion probing to mechanically stable anchoring.

### Prolonged talin binding by RIAM alters adhesion mechanics without recapitulating native KANK function

Based on prior work, particularly Sun et al. (2016), one would predict that engaging talin through a KANK-derived talin-binding sequence would reduce traction force transmission by interfering with actin binding at talin’s ABS2. Native KANK proteins bind talin within the R7–R8 region and recruit cortical microtubule–stabilizing complexes, while simultaneously attenuating force propagation through the integrin–talin–actin axis. Under this model, incorporation of a Kank2 talin-binding motif into RIAM would be expected to suppress traction forces and stabilize adhesions (31). Contrary to this expectation, we observe the opposite phenotype: the RIAM–Kank2TBS chimera enhances traction force generation and accelerates adhesion turnover. This discrepancy argues that disruption of actin binding at ABS2 is unlikely to be the dominant driver of the observed phenotype. Importantly, the effects of the chimera are partially Rap1 dependent, as mutation of the Rap1-binding site (K210E) substantially attenuates adhesion dynamics and force output. If the phenotype were primarily due to steric interference with actin binding at ABS2, Rap1 binding would not be expected to modulate these outcomes. Instead, the Rap1 dependence points to a RIAM-specific regulatory mechanism rather than a generic KANK-like effect on talin–actin coupling.

We propose that the key distinction instead lies in the *temporal layering* of signaling and mechanical reinforcement at talin. Importantly, vinculin recruitment to talin, primarily through unfolding of the R2/R3 domains, can still occur in cells expressing the RIAM–Kank2TBS chimera, enabling normal reinforcement of talin–actin connections via vinculin, as in wild-type adhesions (32, 33). Thus, the elevated traction forces observed in chimera-expressing cells cannot be explained by impaired vinculin binding or weakened actin coupling. Rather, they reflect the *superposition* of persistent RIAM–Rap1 activity on top of an otherwise intact vinculin-mediated mechanical linkage. By stabilizing RIAM–talin engagement, the chimera prolongs Rap1-dependent signaling at adhesion sites, enhancing integrin activation, actin polymerization, and adhesion nucleation even as vinculin-mediated force transmission proceeds. This sustained Rap1–RIAM signaling biases adhesions toward rapid assembly and accelerated turnover, while simultaneously increasing traction output through continued talin engagement and actomyosin coupling. In this framework, vinculin does not replace RIAM functionally but rather reinforces the mechanical axis while RIAM continues to drive a dynamic, signaling-active adhesion state. These observations suggest that timely replacement of RIAM by vinculin is not required to initiate force transmission, but is instead essential to *terminate* Rap1-dependent adhesion dynamics. Failure to remove RIAM prolongs a high-turnover, signaling-dominant state that amplifies traction but destabilizes adhesion lifetime. Thus, vinculin-mediated maturation serves as a critical checkpoint that converts early, Rap1-driven adhesion dynamics into stable, load-bearing focal adhesions.

Our observation that the RIAM-Kank2TBS chimera accelerates FA assembly likely reflects more than just enhanced integrin activation. While replacing RIAM’s autoinhibited TBS1 with the talin-1-binding motif from Kank2 strengthens its integrin-activating capacity, RIAM’s N-terminal domain also binds actin regulators such as profilin and Ena/VASP (8), which promote localized actin polymerization at NAs. This actin linkage supports membrane protrusion and stabilizes new adhesions, contributing to faster assembly. Additionally, the N-terminus of RIAM is phosphorylated by focal adhesion kinase (FAK), and such phosphorylation has been shown to relieve autoinhibition and amplify RIAM’s activity (15). Importantly, FAK itself plays an active role in FA assembly (34) by recruiting talin-1 and promoting pro-assembly signaling through its kinase activity. Together, these mechanisms likely synergize in the RIAM-Kank2TBS chimera, enabling it to coordinate integrin engagement, cytoskeletal remodeling, and signaling amplification—thereby driving accelerated adhesion assembly beyond what integrin activation alone would accomplish.

Our finding that the RIAM-Kank2TBS chimera accelerates FA disassembly reveals a novel role for RIAM in not only assembling but also dismantling adhesions, and highlights that disrupting RIAM autoinhibition can enhance adhesion turnover. Previous studies have shown that RIAM depletion leads to enlarged, stable adhesions with impaired disassembly and increased paxillin phosphorylation, suggesting that RIAM facilitates FA turnover (23). Moreover, FAK, a known regulator of adhesion disassembly (22, 24), can phosphorylate RIAM at Tyr-45 to relieve its autoinhibition (15), potentially enhancing RIAM dynamics during adhesion turnover. By releasing RIAM from its autoinhibited state, the Kank2TBS chimera may allow more rapid recruitment and removal of RIAM at adhesion sites, promoting the disassembly process. Our data also show that Rap1 binding is not essential for disassembling adhesions per se, but contributes to efficient RIAM dissociation, indicating that Rap1 facilitates the recycling or clearance of RIAM as adhesions are disassembled. These results support a model in which the autoinhibited state of RIAM not only controls adhesion assembly but also restrains its participation in disassembly, and that bypassing this inhibition, via forced adhesion targeting, promotes faster adhesion turnover and dynamic cell behavior.

Collectively, our results position RIAM as a molecular timer that links integrin activation to adhesion mechanics through regulated engagement with talin. RIAM activation and removal represent coupled steps that define adhesion lifetime and mechanical behavior. Prolonged RIAM–talin association perturbs this balance, leading to elevated traction forces and rapid turnover, and highlighting the importance of subsequent vinculin recruitment in stabilizing mechanically mature adhesions. Rather than serving solely as an initiator of integrin activation, RIAM emerges as a transient regulator whose residence time shapes the transition from dynamic, force-generating adhesions to stable, load-bearing structures.

## Methods

### cDNA constructs and antibodies

Inducible 3T3 cell line that expresses GFP-RIAM, GFP-RIAM-Kank2TBS or GFP-Kank2TBS-RIAM (K210E) was generated by T-REx^TM^ System in Flp-In™-3T3 cell line (ThermoFischerScientific, Waltham, MA), respectively. Authenticity of constructs was validated by DNA sequencing. In the experiment, doxycycline at 20ng/ml showed no spectral bleed-through, confirming the specificity of the fluorescence signal. Anti-HA monoclonal antibody (Biolegend, San Diego, CA) and anti-GFP rabbit polyclonal antibody (TaKaRa Bio USA, Inc., San Jose, CA) were commercially purchased. Anti-HA mouse monoclonal antibody (12CA5), αIIbβ3 activation-specific mouse monoclonal antibody PAC1, αIIbβ3 activating mouse monoclonal antibody anti-LIBS6 (Ab33) (35), and the αIIbβ3-specific competitive inhibitor integrilin were previously described (9).

### Cell culture and chemicals

3T3 fibroblasts stably expressing paxillin-mRuby and inducible GFP-RIAM, GFP-RIAM-Kank2TBS or GFP-Kank2TBS-RIAM (K210E) and CHO-K1 cells were cultured in 5% CO_2_, 37 °C incubator using Dulbecco’s Modified Eagle Medium (ThermoFisher) supplemented with 10% fetal calf serum, non-essential amino acids and glutamine.

### Live-cell adhesion imaging

3T3 fibroblasts stably expressing paxillin-mRuby and inducible GFP-RIAM, GFP-RIAM-Kank2TBS or GFP-Kank2TBS-RIAM (K210E) were imaged under total internal reflection fluorescence (TIRF) microscope (optoTIRF, CAIRN Research) housed in Nikon Ti-S microscope (Nikon Instruments) at a 60x objective. The microscope stage is equipped with an H301 stage-top incubator chamber and UNO controller (Okolab USA Inc, San Bruno, CA) to maintain cells at 5% CO2 and 37°C in a humid environment. The cells were kept in focus using the CRISP autofocus unit (ASI Applied Scientific Instrumentation, USA). Cell’s GFP was induced by applying doxycycline (20 ng/mL) 20 hours before imaging. Both the mRuby signal and GFP signal of cells cultured on a 35 mm, No. 1.0 Coverslip, 14 mm Glass diameter, glass-bottom dish (MatTek, USA) were imaged on 587nm and 488nm lasers with 500ms exposure time for 60 minutes with 20 seconds frame-to-frame time interval. The frames were captured with a Hamamatsu orca-flash 4.0 LT plus CMOS camera (Hamamatsu Corporation, Bridgewater. NJ, USA) and controlled with MetaMorph imaging software (Molecular Devices, Downingtown, PA).

### Measurement of Integrin αIIbβ3 Activation

Chinese hamster ovary (CHO) cells stably expressing αIIbβ3 were transiently co-transfected with cDNAs encoding HA-talin and either GFP-RIAM, GFP-kank2TBS-RIAM, or GFP alone using Lipofectamine 3000 (Invitrogen). After 24 hours, GFP-positive cells were harvested, stained with PAC-1 antibody, washed, and subsequently labeled with Alexa647-conjugated goat anti-mouse IgM (Invitrogen). Five minutes before flow cytometric analysis, propidium iodide (PI; final concentration 2 μg/ml) was added to exclude dead cells. Cells were analyzed using a BD Accuri™ C6 Plus flow cytometer (BD Biosciences). PAC-1 binding was assessed in live (PI-negative), transfected (GFP-positive) cells. Integrin activation was quantified using the activation index (AI), calculated as: *AI* = (*F* − *F*_0_)/(*F*_*max*_ − *F*_0_), where F is the mean fluorescence intensity (MFI) of PAC-1 binding, F_o_ is the MFI of PAC-1 binding in presence of the αIIbβ3 antagonist integrilin (1 μM), and F_max_ is the MFI of PAC-1 binding in the presence of αβ3-activating mAb anti-LIBS6 (2 μM) (36).

### Focal adhesion analysis

The adhesion analysis was done using the focal adhesion package as previously described (29, 30). Briefly, from images of paxillin-mRuby, NAs and FAs were captured separately. First, NAs were detected using the point source detection method as previously described (37). Then, FAs and focal complexes (FCs) were segmented using thresholding, by combination of Otsu and Rosin, the paxillin images processed with Gaussian filtering and background subtraction. Segmented areas were categorized using criteria (38) in which areas larger than 0.2 µm^2^ were considered either for FCs or FAs (0.24 µm^2^ for FCs and 0.6 µm^2^ for FAs). The length and width of the individual segmentations were evaluated by length of the major axis and width of the minor axis in an ellipse respectively that fit in each segmentation. The FA eccentricity was assessed by the ratio of the length of the minor axis over the length of the major axis in the ellipse fit to each segmented FA or FC. Nascent adhesions (NAs) were detected using point source detection method as previously described (37). Briefly, paxillin and GFP images were filtered employing the Laplacian of Gaussian filter and then local maxima was detected. Each local maximum was then fitted with an isotropic Gaussian function (standard deviation: 2.1 pixels, i.e., ∼180 nm) and outliers were removed using a goodness of fit test (p= 0.05). The point sources detected for NAs were tracked over the entire frames of the time-lapse images using our Focal Adhesion Package (30) and compared with FC/FA segmentations to determine their maturation state. Lifetimes of adhesions were calculated from the lifetimes of individual tracked trajectories. The lifetimes of NAs and FAs were separately assessed. To do that, each trajectory has stored a field called ‘status’, i.e., ‘BA (before adhesion)’, ‘NA’, ‘FC’, ‘FA’, etc., depending on its detection or overlap with segmentation. Whereas NA lifetimes were counted from trajectories of NAs, i.e., no overlap with FC/FA segmentation, lifetimes of FCs/FAs were counted from trajectories that contain FC or FA status at any time in their lifetimes.

### Assembly and disassembly rate analysis

The rate of assembly and disassembly was determined by measuring the fluorescent intensity of individual adhesions from paxillin and GFP images over time, with the background fluorescent intensity subtracted. Rate constants for assembly and disassembly were determined from the slope of the graphs Log (I/I_0_) and Log (I_0_/I), respectively, over time, in which I_0_ is the initial fluorescent intensity and I is the fluorescent intensity at the indicated times (39). To detect the most appropriate period for each rate constant determination, the adjusted R^2^ values from the linear models for various time periods were assessed, and the model with the largest adjusted R^2^ value was selected, from which the slope was recorded (40). The multiple linear models for the assembly rate were determined by setting the intensity at the very initial time point as I_0_ and varying the later time points until the maximum intensity. The models for the disassembly rate were determined by varying the time points that are earlier than the last time point until the maximum intensity. The intensity at the earliest time point of the best linear model was chosen as I_0_. The best adjust R^2^ value per adhesion was again used to filter out noise-like trajectories, for which we used 0.3 as the threshold.

### Traction force imaging and analysis

TFM experiment was done on 5 kPa silicone gel as previously (29, 41). Briefly, the TFM gel substrate was fabricated by spin-coating a 35 mm glass-bottom dish (#1 cover glass, from MatTek corporation) with a high-refractive-index silicone gel. To visualize the gel deformation, carboxylated far-red fluorescent beads (excitation/emission 690/720 nm, from Invitrogen) with 40 nm diameter were covalently bonded on the gel surface with a density of 1 bead /mm2 using EDC chemistry. For imaging, the 3T3 cells expressing inducible GFP-RIAM, GFP-Kank2TBS-RIAM, or GFP-Kank2TBS-RIAM (K210E) were cultured in Dulbecco’s Modified Eagle Medium without phenol red (Gibco, 31053) supplemented with 10% fetal bovine serum, 1% penicillin/streptomycin, 1% nonessential amino acids, and 1% L-glutamine. A single-shot live cell imaging was performed using the same TIRF microscope setup on 642 nm (beads) with 800nm (GFP-RIAM & GFP-Kank2TBS-RIAM) and 500 nm (GFP-Kank2TBS-RIAM (K210E)), 587 nm (paxillin) and 488 nm (GFP) with 500 nm exposure time. The bead images with relaxed gel were obtained after removing the cells using 0.5 ml of 10% bleach.

Traction force reconstruction was done as previously described (29). Briefly, using a pair of bead images acquired in the presence and absence of cells, the displacement field was calculated by particle tracking velocimetry method, which is an image cross-correlation-based tracking method after identifying centers of all individual beads in the field of view. The template size of 17 pixels and a maximum displacement of 50 pixels were used for defining an interrogation area. The displacement field was corrected using outlier filtering. The force reconstruction was performed using Fast Boundary Element Method (FastBEM) with L2-norm-based regularization, where a regularization parameter was chosen using the L-curve, L-corner method (29).

## Author contributions

H.S.L: conceptualization, data curation, formal analysis, investigation, methodology, validation, writing, and editing. N.M: conceptualization, data curation, formal analysis, investigation, methodology, validation, writing, and editing. S.J.H: conceptualization, formal analysis, funding acquisition, methodology, project administration, supervision, visualization, and writing—original draft, review, and editing. M.H.G: conceptualization, formal analysis, funding acquisition, methodology development, project management, oversight, visualization, and writing—including the original draft, review, and editing.

## Conflict of Interest

The authors declare no competing interests.

## Acknowledgment

This work was supported by a NIH R15GM135806 grant (to S.J.H.) and NIH PO1HL151433 (to M.H.G.). N.M. would also like to thank Health Research Institute (HRI) at Michigan Technological University for the HRI graduate student fellowship. N.M. is also grateful to Dr. Michael Fleming and Dr. Susan Skochelak in providing Fleming & Skochelak Graduate Fellowship in Human Health to support his research.

## Supplementary Figures

**Figure S1.**
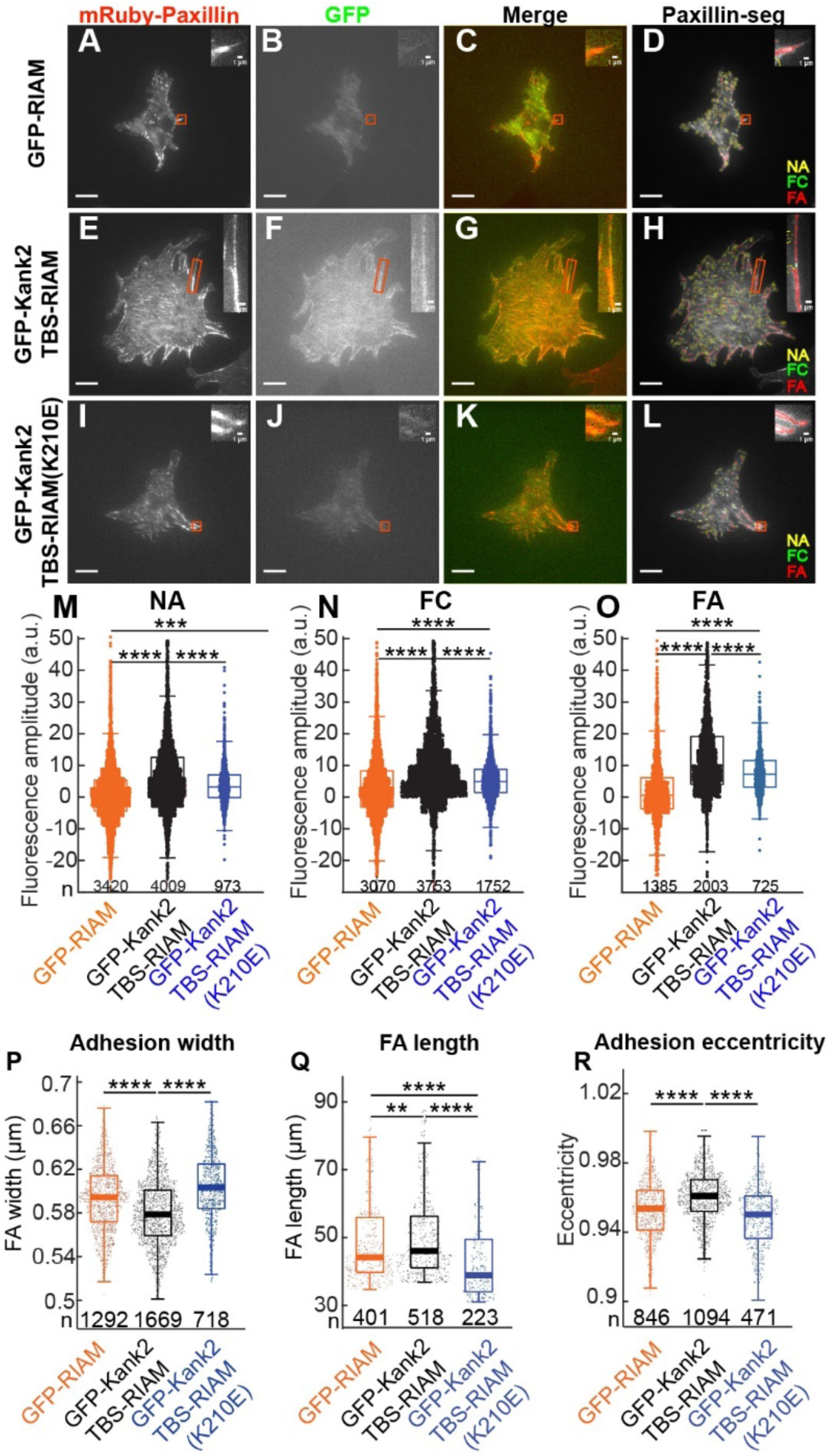
RIAM-Kank2TBS colocalizes with FA in a size-dependent manner and makes adhesions longer and thinner in a Rap1-dependent manner. (A-L) TIRF images of mRuby-paxillin **(A,E,I)**, GFP **(B,F,J)**, merge **(C,G,K)** and paxillin images with adhesion segmentation **(D,H,L)** of representative 3T3 fibroblasts expressing mRuby-paxillin and GFP-RIAM **(A-D)**, mRuby-paxillin GFP-RIAM Kank2TBS **(E-H)**, and mRuby-paxillin GFP-RIAM Kank2TBS (K210E) **(I-L)** on a 5kPa gel substrate. Scale bar: 20 μm. GFP-tagged RIAM and its variants were induced by 20 ng/ml of doxycycline. Adhesion segmentation shows NAs in yellow, focal complexes in green, and FAs in red. Inset: zoomed-in images displaying a representative FA in each red box. Inset scale bar: 1 μm; adhesion segmentation in FA (green), focal complex (red), NA (yellow). **(M-O)** Box plots of fluorescence intensity of NA (**M**), focal complex (FC, **N**) and FA (**O**) in GFP channel. **(P-R)** Box plot of FA width **(P)**, FA length **(Q)**, FA eccentricity **(R)**. The numbers shown on top of the x-axis are the numbers of adhesions collected from independently imaged cells, for GFP-RIAM (n=30), n=30 for GFP-RIAM Kank2TBS, and n=16 for GFP-RIAM Kank2TBS (K210E). **: p<1×10^-3^, ***.: p<1×10^-5^, ****.: p<1×10^-10^ by Mann-Whitney U test.

## Notes

### Competing Interest Statement

The authors have declared no competing interest.

